# Prospective evaluation of beta-lactamase detection in penicillin susceptible *Staphylococcus aureus* by interpretation of the penicillin disc edge

**DOI:** 10.1101/776880

**Authors:** Andrew Henderson, David Whiley, Fleur Francis, Cathy Engler, Matthew Glover, Joel Douglas, Letitia Gore, Patrick Harris, David Paterson, Graeme Nimmo, Robert Norton

## Abstract

Penicillin susceptible *Staphylococcus aureus* (PSSA) may occasionally be encountered as a cause of complicated *S. aureus* infection, such as endocarditis or bloodstream infections. Clinicians may choose to treat these patients with penicillin over a semi-synthetic penicillin derivative, such as flucloxacillin or oxacillin, due to a favourable Pk/Pd profile. In this study, we prospectively evaluated the penicillin disc (1-IU) method for detection of *blaZ*, with interpretation of the penicillin edge according to EUCAST recommendations. 472 PSSA isolates were collected between September 2014 to December 2015 from three clinical microbiology laboratories in Queensland, Australia. Initial antimicrobial susceptibility testing was performed by the Vitek 2 system. Real-time PCR for *blaZ* was performed following phenotypic testing with the 1-IU penicillin disc and the PCR used as the gold standard for detection of penicillinase. The prevalence of *blaZ* amongst the isolates was 7%. The sensitivity, specificity, positive predictive value and negative predictive value of the penicillin disc method was 97%, 95%, 61% and 100% when compared to *blaZ* PCR. In summary, the penicillin disc zone size and edge interpretation is a reliable method for detection of *blaZ* in *S. aureus* isolates that otherwise test susceptible to penicillin by Vitek 2 AST.

## Introduction

Penicillin is commonly used to treat patients with complicated infections with invasive penicillin susceptible *Staphylococcus aureus* (PSSA) infections.(1) Key advantages of penicillin treatment include a lower MIC distribution compared with other beta-lactam agents active against *S. aureus*, as well as the ability to achieve higher free non-protein-bound plasma drug concentrations.(1) However, the recently updated American Heart Association guidelines on infective endocarditis do not recommend the use of penicillin for PSSA endocarditis.(2) This decision is primarily based on concerns raised over the ability of laboratory methods to accurately detect certain forms of penicillin resistance in *S. aureus*; specifically, current evidence indicates that screening methods for the *blaZ* penicillinase gene are insensitive.(2)

Two key mechanisms are responsible for resistance to penicillin in staphylococci; the *blaZ* gene which encodes for penicillinase, a serine beta-lactamase which hydrolyses the β-lactam ring resulting in the production of penicilloic acid, and the *mecA* gene which encodes for penicillin-binding protein (PBP) 2A.(3, 4) While the presence of *mecA* can readily and accurately be detected through the use of the cefoxitin disk diffusion test, there are well-recognized difficulties with detection of *blaZ*.(5) The nitrocefin hydrolysis test, previously a widely used phenotypic method for penicillinase detection amongst *S. aureus*, has been shown to produce false negative results.(6-10) Molecular methods (including PCR) may be more be accurate, but are typically too expensive and slow compared to the phenotypic methods to be a suitable alternative for routine use. Molecular assays may also be impeded by the nucleotide variations in sequence targets associated with the different *blaZ* types.(10) At present, both CLSI and EUCAST recommend phenotypic testing via a penicillin disc.(11) By utilizing the penicillin disc method, isolates harbouring *blaZ* may still appear sensitive according to the zone size, but *blaZ* presence may be identified by closer inspection of the zone edge, and is indicated by the appearance of a straight edge or cliff. Although being the current recommended method, limited prospective data is available to confirm the validity of penicillin disc detection of *blaZ* in clinical microbiology laboratories. In fact, experiences at our laboratory suggest that scientists, particularly those infrequently performing the method, struggle to appropriately interpret the zone edges.

The study was designed to address two aims. The first aim was to use PCR to examine the prevalence of the *blaZ* beta-lactamase amongst PSSA isolates across 3 microbiology laboratories in Queensland. The second aim was to prospectively evaluate the 1 IU penicillin disc test for detection of penicillinase in PSSA isolates compared to the nitrocefin test, using real-time PCR for *blaZ* as the gold standard.

## Methods

### Overview

This project was performed in a clinical microbiology laboratory in the city of Townsville, located in the state of Queensland, Australia. From September 2014 to December 2015, isolates (total = 472) were prospectively collected for the study from 3 laboratories (Townsville Hospital, Gold Coast Hospital, Princess Alexandra Hospital).

For isolates collected in Townsville (n = 207), the interpretation and report of the penicillin zone size and edge was performed by scientific staff as part of the routine standard operating procedure; note that this zone test was the sole method used for routine detection of *blaZ* when the Vitek 2 demonstrated penicillin susceptibility. Isolates were then stored and later batch-tested as part of the research study using the nitrocefin test and *blaZ* PCR and retested using the penicillin disc test by the investigators (supervising scientist and microbiology registrar). Isolates collected in the Gold Coast Hospital (n = 65) and Princess Alexandra Hospital (n = 200) laboratories were stored and later transferred and batch-tested as above (nitrocefin, *blaZ* PCR and penicillin disc tests) by the investigators at the Townsville laboratory. It should be noted that all penicillin disc tests were performed independently of and without knowledge of the nitrocefin or *blaZ* PCR test results. Where isolates demonstrated discrepant results between *blaZ* PCR and the penicillin disc test or nitrocefin results, the isolate was repeated by all 3 tests and interpreted by two investigators blinded to each other’s interpretation.

### S. aureus identification and susceptibility testing

Identification of *S. aureus* was performed at all three laboratories via a combination of latex agglutination for coagulase activity, presence of DNase (DNase test agar, Oxoid, Thermo Fisher Scientific), Vitek 2 (bioMerrieux) identification and MALDI-TOF mass spectrometry (bioMerrieux). All *S. aureus* isolates underwent antimicrobial susceptibility testing, performed with the Vitek 2 automated broth microdilution antimicrobial susceptibility testing (AST) system, with susceptibility interpreted against EUCAST criteria.(11) Only isolates with penicillin MICs ≤ 0.125 μg/ml (EUCAST penicillin breakpoint) were included in the study.

### Penicillin disc test

A 0.5 MacFarland suspension was made from a single colony of *S. aureus* for Kirby Bauer disc diffusion test on Mueller-Hinton agar (MHA, bioMerrieux), with a 30 μg cefoxitin disc and 1 μg penicillin disc incubated at 35 degrees Celsius in atmospheric oxygen for 24 hours. Zone sizes were measured and interpreted according to EUCAST criteria. Isolates with a zone < 26 mm, or with a sharp zone edge were considered resistant.(11)

### Nitrocefin test

Phenotypic beta-lactamase activity was performed on all isolates using nitrocefin (SR12C; Oxoid) by the paper disc spot method, according to the manufacturer’s instructions. A sweep of colonies were taken from the edge of the cefoxitin disc, which was incubated for 24 hours at 35 degrees Celsius, and applied to the nitrocefin impregnated region. A change in colour of the paper from yellow to red indicated a positive result and this was read after 15 minutes, 30 minutes and 1 hour.

### blaZ PCR

Bacterial DNA was extracted from pure colonies growing on Columbia horse blood agar by the boiling lysis method as follows. Two to three colonies of *S. aureus* were suspended in 0.5ml of sterile demineralised water and immersed in a water bath at 100 degrees Celsius for 10 minutes. After this, the suspension was centrifuged at 15000 x g for 30 seconds. 50 µl of supernatant was stored at - 80 degrees Celsius, in preparation for performing nucleic acid amplification testing (NAAT).

Real-time PCR amplification of the *blaZ* gene was performed with the previously described primers and Taqman probe by Pereira, *et al*. (10), with some modifications. Briefly, the reaction mix contained 0.5 µM forward primer (5’-GCT TTA GAA CTT ATT GAG GCT TCA-3’), 0.5 µM reverse primer (5’-CCA CCG ATY TCK TTT ATA ATT T-3’), 0.2 µM Taqman probe (5’-FAM-AGT GAT AAT ACA GCA AAC AA-MGBNFQ-3’, where FAM is 6-carboxyfluoroscein), 12.5 µl QuantiTect Probe PCR buffer mix (Qiagen, Australia) and 5 µl of nucleic acid extract in a total volume of 25 µl per test. The primers and probe were sourced from Integrated DNA Technologies. Amplification was performed using RotorgeneQ real-time thermocyclers (Qiagen, Australia), with reactions run under the following conditions: 95 °C for 10 minutes, followed by 45 cycles of 95 °C for 15 seconds and 60 °C for 60 seconds. Fluorescent probe signals were read during the 60 second extension step of the cycling program.

### Statistical analysis

All statistical analysis was performed using R (R Project for Statistical Computing, http://www.r-project.org/). Graphs were created though the ggplot2 package with the exception of the receiver operator curve for zone size and *blaZ* detection, which was created through the pROC package.(12) Cohen’s Kappa score was used to determine the agreement between scientific staff and investigators for interpretation of the penicillin zone edge for the isolates collected in Townsville. McNemar’s test was performed to compare sensitivity and specificity, whereas weighted generalized scores were performed to compare positive and negative predictive values for paired isolates (13, 14). A p-value <0.05 was considered statistically significant.

## Results

The overall prevalence rate of *blaZ* amongst the combined 472 PSSA isolates was 7.3% (n = 34, 95% confidence interval 5.3% - 10.1%) according to the *blaZ* PCR results. This result was similar amongst the 207 isolates from the Townsville Hospital laboratory (prevalence = 6.8%, 95% CI 4.0 - 11.0%), the 65 isolates from the Gold Coast Hospital laboratory (10.8%, 95% CI 0.8% - 13.2%) and the 200 isolates from the PA Hospital laboratory (6.5%, 95% CI 3.8% - 10.6%).

Table 1 demonstrates the sensitivity, specificity and predictive values calculated for the penicillin disc test and nitrocefin test for *blaZ* detection (using the *blaz* PCR as the gold standard). In comparison to the penicillin disc test, nitrocefin performed poorly at detecting *blaZ* amongst PSSA isolates. The sensitivity of nitrocefin (15%, 95% CI 5 – 31%) was lower than the penicillin disc test (97%, 95% CI 85 – 100%) for detection of *blaZ*, with the difference in sensitivity between the penicillin disc test and nitrocefin found to be statistically significant (p < 0.01). Despite the low prevalence rate of *blaZ* amongst the isolates, a statistically significant difference between the negative predictive value (NPV) of nitrocefin and the penicillin disc test was detected (94% versus 100%; p <0.01)

**Table 1.** Sensitivity, specificity, negative and positive predictive value of the penicillin disc test in comparison to nitrocefin.

|  | Penicillin 1-IU Disc Test |  |  |  | Nitrocefin<sup>α</sup><br>(n = 472)<br>% (95% CI) | P Value <sup>β</sup> |
| --- | --- | --- | --- | --- | --- | --- |
|  | Townsville Hospital Isolates (n = 207) |  |  | Overall<sup>α</sup><br>(n = 472)<br>% (95% CI) |  |  |
|  | Scientists<br>% (95% CI) | Investigators<br>% (95% CI) | P Value |  |  |  |
| Sensitivity<sup>a</sup> | 93 (79 – 100) | 100 (100 – 100) | 1 | 100 (100 – 100) | 15 (5 – 31) | <0.01 |
| Specificity<sup>b</sup> | 93 (90 – 97) | 98 (97 – 100) | 0.01 | 97 (96 – 99) | 100 (99 – 100) | <0.01 |
| PPV<sup>c</sup> | 50 (31 – 69) | 82 (64 – 100) | <0.01 | 76 (63 – 88) | 100 (36 – 100) | 0.07 |
| NPV<sup>d</sup> | 99 (98 – 100) | 100 (100 – 100) | 0.3 | 100 (100 – 100) | 94 (91 – 96) | <0.01 |

a Sensitivity = True positives/(False negatives + True positives)
b Specificity = True negatives/(False positives + True negatives)
c Positive predictive value = True positives/(False positives + True positives)
d Negative predictive values = True negative/(False negatives + True negatives)
- 376 $\alpha$ Test performed by investigators; includes Townsville laboratory isolates, Gold Coast laboratory isolates and Princess Alexandra Hospital isolates. - 378 $\beta$ Comparison between penicillin disc and nitrocefin

Figure 1 demonstrates a histogram of penicillin zone size for the isolates based upon detection or absence of *blaZ*. A receiver operator curve (Supplementary Figure 1) was derived from the zone sizes to determine sensitivity and specificity of zone size for detection of *blaZ* and is demonstrated in Supplementary Table 2. For isolates with a Vitek 2 MIC of ≤ 0.03 μg/ml, *blaZ* was detected in 4/83 isolates, compared to 8/275 and 22/114 with MICs of 0.06 μg/ml and 0.12 μg/ml respectively.

**Figure 1.**
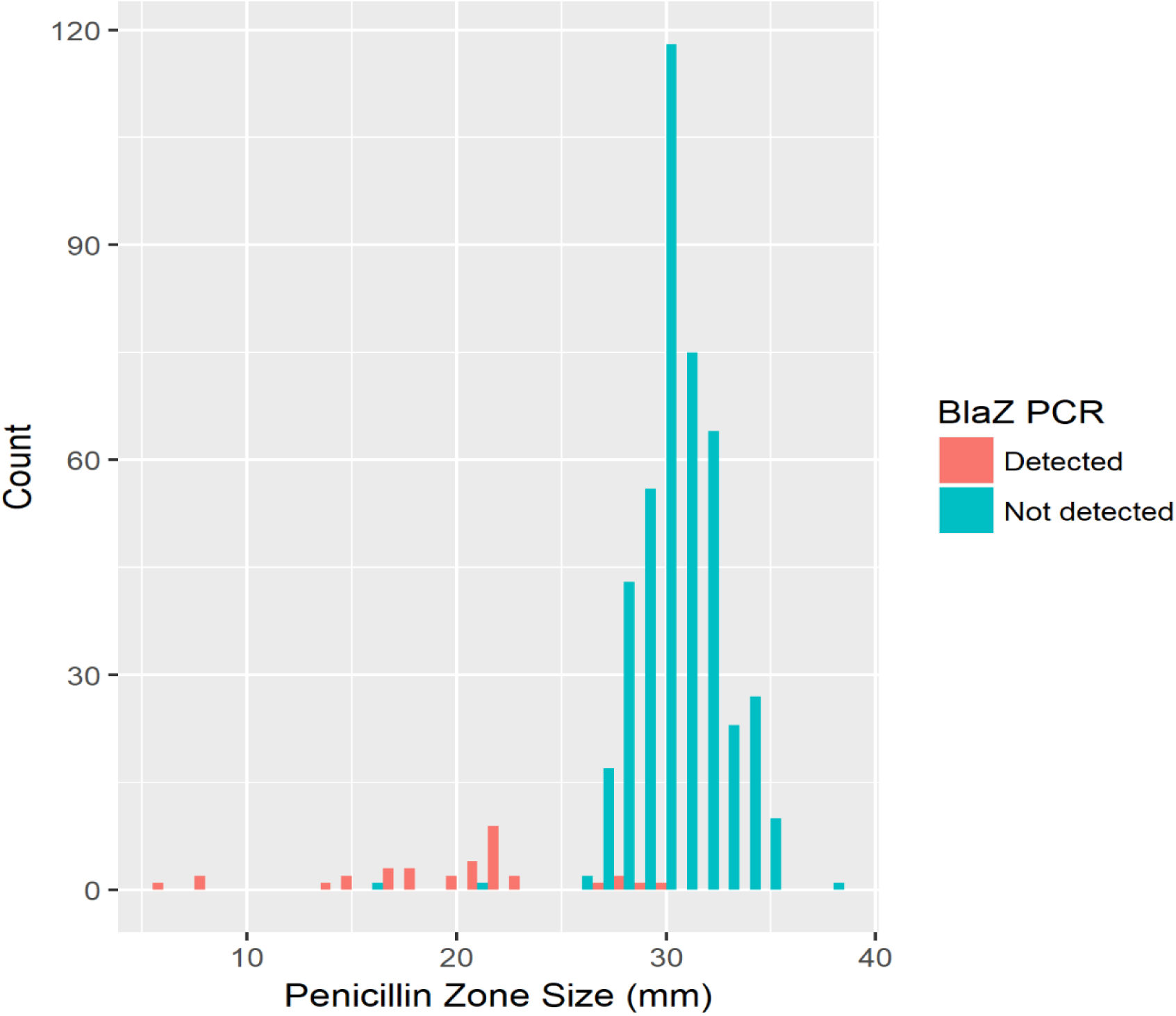
Histogram of Penicillin Zone Size According to Presence of *blaZ*.

Two isolates were found to have penicillin zone sizes less than 26mm (EUCAST breakpoint) yet *blaZ* was not detected by PCR. Both isolates demonstrated sub-populations of growth within an outer zone of inhibition to the penicillin disc that measured greater than 26 mm. The edge of the outer zone appeared fuzzy, indicating absence of beta-lactamase. In addition, both isolates (inner and outer populations) tested negative for penicillinase activity by nitrocefin. The sub-population colonies growing inside the outer zone were confirmed to be *S. aureus* when identification was repeated. Vitek 2 AST was then performed on the sub-population colonies growing within the zone of inhibition with results now demonstrating penicillin MICs ≥ 0.5 μg/ml, yet both isolates were now found to have oxacillin MICs ≥ 2 μg/ml and were cefoxitin screen positive. The sub-population colonies demonstrated heavy growth on MRSA chromogenic agar (Biomerrieux). To confirm the presence of *mecA*, Xpert MRSA/SA assay (Cepheid, Sunnyvale, CA) was performed on both the outer zone and inner zone populations. The inner zone population was confirmed as MRSA with *mecA* detected, however the outer population did not have *mecA* detected.

Overall, 25 isolates produced initial discordant phenotypic results when compared to *blaZ* PCR, with all discordant results resolved upon repeat testing by the investigators (Supplementary Table 2. 24 *blaZ* negative isolates were incorrectly reported as resistant (13 by scientists and 11 by investigators) and 1 *blaZ* positive isolate was incorrectly reported by a scientist as susceptible to penicillin when interpreting the penicillin disc test. Agreement by Cohen’s Kappa score between scientists and investigators for Townsville isolates was 0.58 (moderate agreement).

## Discussion

In this study, we investigated beta-lactamase detection amongst PSSA isolates in a routine clinical microbiology laboratory following the recent introduction of the penicillin disc test to replace the nitrocefin test. The results of our study, along with other recently reported studies, confirm that the penicillin disc test (using EUCAST methodology) is a reliable predictor of phenotypic activity of *blaZ*.(8-10) Notably, only 1 isolate that was positive for *blaZ* by PCR was not detected by the penicillin disc test. This isolate was tested by scientists as part of routine practice and was incorrectly reported as susceptible to penicillin. The isolate was subsequently confirmed as penicillin resistant by the investigators when the test was repeated and the discrepancy was considered to be caused by human error, rather than a problem with the penicillin disc test. Consistent with previous studies, we also found the nitrocefin test to be highly insensitive and re-enforces the decision of our local laboratories to move away from the nitrocefin test as per the updated EUCAST recommendations.(6, 8, 9)

Overall the prevalence rate of *blaZ* detected in PSSA from the three laboratories was 7%. This rate is lower than from recently reported SAB data in 2013, in the Australian setting, where 332 (20.4%) isolates initially tested susceptible to penicillin from 1342 episodes of MSSA bacteraemias and beta-lactamase was detected in 69 (20.8%) of the PSSA isolates.(15) These rates support the recommendation made by EUCAST to screen for *blaZ* amongst PSSA in our region prior to reporting isolates as susceptible to penicillin.(11) Although there are theoretical advantages to using penicillin over other agents for PSSA infections, the use of penicillin for this indication needs to be considered against the capacity of the laboratory to detect *blaZ*. Importantly, the NPV of the penicillin disc test was 99% when performed by the routine scientists and 100% by the investigators. These high NPVs suggest that if clinicians were to treat patients with penicillin on the basis of a negative penicillin disc test result then the use of use penicillin would be appropriate in at least 99% of cases. In contrast, the significantly lower NPV of the nitrocefin test (94%, p < 0.01) would result in approximately 6% of patients (infected with *blaZ*-harbouring strains) inappropriately receiving penicillin treatment. Although the positive predictive value (PPV) of the disc test was lower than the nitrocefin method (76% vs 100%), this issue (reporting of false resistance) was not considered a major limitation of the disc test as it would only limit treatment options, rather than leading to incorrect treatment. Also, clinicians may otherwise choose to use flucloxacillin over penicillin regardless of the laboratory reporting penicillin as susceptible.(16) As such, it is our opinion that the potential benefit of the PPV of nitrocefin should not be relied upon when laboratories make a decision in regards to the choice of test for *blaZ* detection.

An unusual finding in this study was the identification of 2 subpopulations of MRSA that were *blaZ* negative. Heterogenous expression of methicillin resistance was able to be identified by induction of sub-populations expressing resistance through prolonged beta-lactam exposure using penicillin discs. A likely explanation for this is the presence of two distinct *S. aureus* populations (hetero-resistance), one carrying the *mecA* gene and one without, but with both populations negative for *blaZ*. It is possible that *mecA* expression was altered in the absence of *blaZ* given that *mecA* induction occurs at a much slower rate than that of *blaZ* induction, with a recent study demonstrating that expression of *mecA* relies upon an intact *blaZ* system.(17) Optimal expression of methicillin resistance, occurring when both regulatory systems are present, is due to the formation of BlaI:MecI heterodimers that weakly bind to the *mecA* operon and are then more susceptible to BlaR1 preoteolysis and inactivation, thereby resulting in transcription of *mecA*.(17)

Although our study demonstrated a reliable method for phenotypic detection of beta-lactamase in PSSA isolates there are some limitations. Firstly, we cannot exclude that *blaZ* variants may not have been detected by the primers and probe used in this study. Similarly, we did not perform any sequencing of the real-time PCR products to confirm the specificity of the assay. In addition, many laboratories only perform beta-lactamase testing in PSSA on isolates from blood culture, or sterile sites, where a long phase of intravenous and/or oral antibiotics may be required. Therefore, for laboratories that perform the test on an ad-hoc basis, accurately discerning the edge of the penicillin disc may prove more challenging. This may explain why the specificity result was shown to be lower for scientists, than for the investigators (93% versus 97%). This point is reinforced in the EUCAST guidelines for clinical breakpoints.(11) A clear instruction is given to err on the side of caution and to report isolates as resistant if the result of the test is not clear to the observer.(11)

In summary, for clinicians to confidently use penicillin for the treatment of PSSA, we recommend laboratories consider the penicillin disc test over nitrocefin. Notwithstanding this, laboratories performing the penicillin disc test on an ad-hoc basis run the risk of reduced sensitivity and specificity given interpretation of the disc edge may be dependent on the experience of the scientist.

## Acknowledgements

Funding for this study was provided for by a Study Education and Research Committee grant from Pathology Queensland. The authors would like to thank the 3 laboratories from the Gold Coast Hospital, Townsville Hospital and Princess Alexandra Hospital or providing the isolates for the study.

